# “Multimorbidity states with high sepsis-related deaths: a data-driven analysis in critical care”

**DOI:** 10.1101/491712

**Authors:** Zsolt Zador, Alexander Landry, Michael D. Cusimano, Nophar Geifman

## Abstract

Sepsis remains a complex medical problem and a major challenge in healthcare. Diagnostics and outcome predictions are focused on physiological parameters with less consideration given to patients’ medical background. Given the aging population, not only are diseases becoming increasingly prevalent but occur more frequently in combinations (“multimorbidity”). Thus, it is imperative we incorporate morbidity state in our healthcare models.

We investigate effects of multimorbidity on the occurrence of sepsis and associated mortality in critical care (CC) through analysis of 36390 patients from the open source Medical Information Mart for Intensive Care III (MIMIC III) dataset. Morbidities were defined based on Elixhauser categories, a well-established scheme distinguishing 30 classes of chronic diseases. Using latent class analysis (LCA) we identified six clinically distinct subgroups based on demographics, admission type and morbidity compositions. Subgroup of middle-aged patients with health consequences of drug and alcohol addiction had the highest mortality rate, over 2-fold greater compared to other groups with older patients and complex multimorbid patterns. The findings promote incorporation of multimorbidity in healthcare models and the shift away from current single-disease paradigm in clinical practice, training and trial design.

## Introduction

Sepsis remains one of the most serious medical conditions with high mortality and poor prognosis. It is responsible for more than half the in-hospital deaths and it is the most costly disease in healthcare constituting $20.3 billion or 5.2% of all hospitalization expenses (1). Generally, risk assessment scores of mortality in critical care, such as the Simplified Acute Physiology Score (2) (SAPS II), Sepsis related organ failure score (SOFA) (3) or the Oxford Acute Severity Illness Score (4) (OASIS), focus on inpatient physiological data within 24 hours of admission. Only SAPS II incorporates some rare pre-existing chronic conditions (4). Epidemiological studies however, demonstrate substantial effect from underlying diseases, almost doubling mortality in sepsis (5–7). Given these epidemiological findings it is highly relevant to consider pre-existing morbidity states in assessing sepsis related mortality risk. Furthermore incorporating into research initiatives has the potential to make a significant impact on the health care delivery and health outcomes of patients.

Modern healthcare is extending life expectancy, translating into an increasingly aging population where chronic conditions are prevalent. Suffering from more than one illness is termed multimorbidity, and is associated with increased healthcare use, decreased quality of life and higher mortality (8, 9). A population cross section study of 1,751,841 patients in the United Kingdom showed 23.2 % of the entire population suffered from two or more chronic condition (9). The Canadian Community Health Survey (based on the entire population irrespective of whether they had registered with healthcare providers) revealed a prevalence of 12.9% (8) and other research reveals that among Medicare users ages 65-69, the proportion of individuals presenting with multiple diseases increases to 54% (10).

The majority of current medical education, treatment protocols and clinical trial designs are based on the premise that patients have one disease, failing to consider the reality in which most patients suffer with a number of conditions (8, 9, 11). Further, the majority of drug candidates are identified in animal experiments under standardized conditions, homogenous treatment and control groups. The subsequent clinical trial testing then aims to reproduce the same homogeneity in the recruited patients by excluding morbidity (12, 13). We believe that the inability to account for the heterogeneity introduced by multimorbidity is at least in part why a large number of clinical trials fail. There has however been some shift in focus as the identification of patient groups with similar clinical profiles, termed “endophenotypes”, has led to changes in the management of heterogeneous conditions including sepsis (14), asthma (15), and acute respiratory distress syndrome (16).

Critical care is one of the most data-intense health care environments (17) yielding a broad range of high dimensional data (18). This lends itself well to advanced analytics such as machine learning (19, 20), which allows detection of complex, clinically relevant patterns. Latent class analysis has become increasingly utilized in the discovery of clinically relevant patient subclasses (14, 15, 21). This method assumes the existence of several unobserved groups within the data which share clinical properties that are mutually exclusive between groups. We hypothesized the existence of clinically homogenous patient subgroups or endophenotypes in critical care, and these share clinical characteristics, risk profile, and will ultimately facilitate diagnosis and anticipatory management. In this work, we identify several endophenotypes exhibiting very different risk profiles for adverse health outcomes, such as organ failure, sepsis and associated mortality.

## Methods

### Database

We used the third edition of the Medical Information Mart in Critical Care (MIMIC3) database for our analysis. This is a single center database containing longitudinal data on 38,597 adult patients in critical care with 53,423 distinct admissions. Further details on the database are included in supplementary material.

### Definition of morbidities

The MIMIC3 database includes more than 15,693 distinct diagnoses, which are categorized by ICD9 and ICD 10 codes. For a more compact representation of chronic conditions we summarized diseases using the 30 Elixhauser categories (22) based on an algorithm provided by the authors describing the MIMIC3 database (23). The Elixhauser categories are well established to reflect chronic diseases and they have been validated for both ICD9 and ICD 10 (24).

### Clustering and latent class analysis

We performed a preliminary analysis on similarities between diseases based on disease prevalence in the population using k-means clustering. We first computed disease prevalence (proportion of patients affected by the disease) in each age bracket and used Euclidean distance as similarity measure to define cluster similarities.

For the subsequent analysis of detecting subgroups of patients we used latent class analysis (LCA). This technique assumes the existence of unobserved (“latent”) classes within the study group and identifies them by fitting a set of mixture models to the data. In our analysis we followed the methodological steps of determining and verifying latent classes as summarized by Zho et al. We firstly detected the optimal number of classes based on a combination of Bayesian information criteria, Akaike Information criteria, class size and class interpretability. Characteristics of the latent classes were compared using chi square test for categorical and one-way ANOVA for continuous variables. Residual diagnostics were used to verify that the assumptions for ANOVA were not violated; expected values were calculated to verify the assumptions for Chi-Square test were not violated. We also confirmed the preferred choices between classes using logistic regression models assessed using area under the receiver operator curve (AUC’s). Methodological details are described in supplementary material.

### Network visualization

The complex relationships between morbidities were visualized using network analysis, which demonstrates associations that are otherwise difficult to appreciate. In our initial approach when analyzing the entire critical care dataset, network nodes represented variables and the co-occurrence of variables was assessed using a relative risk (RR) measure described previously (25), which essentially represents the risk of co-occurrence for two diseases. Associations over a significance threshold of p<0.05 were included in the network. The metrics of RR is reflected by edge width and disease prevalence is depicted by diameter of the nodes. In the characterization of endophenotype classes, edges were weighted by the number of patients with the disease pair normalized to the total number of patients within the class. We used this measure as RR has been described to underestimate co-occurrences for highly prevalent diseases in the endophenotype classes (25).

## Results

### Heterogeneous morbidity profile in the critically ill

Cohort demographics are summarized in supplementary table S1. From a population of 36,390 patients, 83.68% of admissions were due to an emergency. There was slightly greater proportion of males (57.85%) than females. Two or more medical condition were reported by 77.28% the patients and nearly half of the entire study population were seniors (age 65 or over). The overall prevalence of sepsis was 37.3% (CI95%: 36.7 – 37.9%), the organ failure rate was 37.5% (CI95%: 37-38%) and overall mortality was 10.9% (CI95%: 10.6 – 11.3%). Mortality associated with sepsis and organ failure were 22.1% (CI95%: 20.3-22%) and 18.4% (CI95%: 17.8-19.1%) respectively.

The proportion of patients with multimorbidity increased steadily with age as expected (8, 9) (Fig 1A). However we found that the prevalence of individual disease groups (reflected by Elixhauser categories) had distinct patterns over age brackets (Fig 1B). Specifically three distinct patterns were observed: two showed increasing prevalence with age, whereas the third had peak prevalence in the 25-44 and 45-64 age bracket but lower prevalence in older age groups. Network discovery showed a variety of disease co-occurrences over the cohort suggesting frequent associations between cardiovascular with pulmonary conditions, diabetes, renal failure and hypertension and the co-occurrence of alcohol/drug abuse, liver failure and coagulopathy (Fig 1C). These findings prompted further investigation for patient subgroups within our cohort.

**Figure 1:**
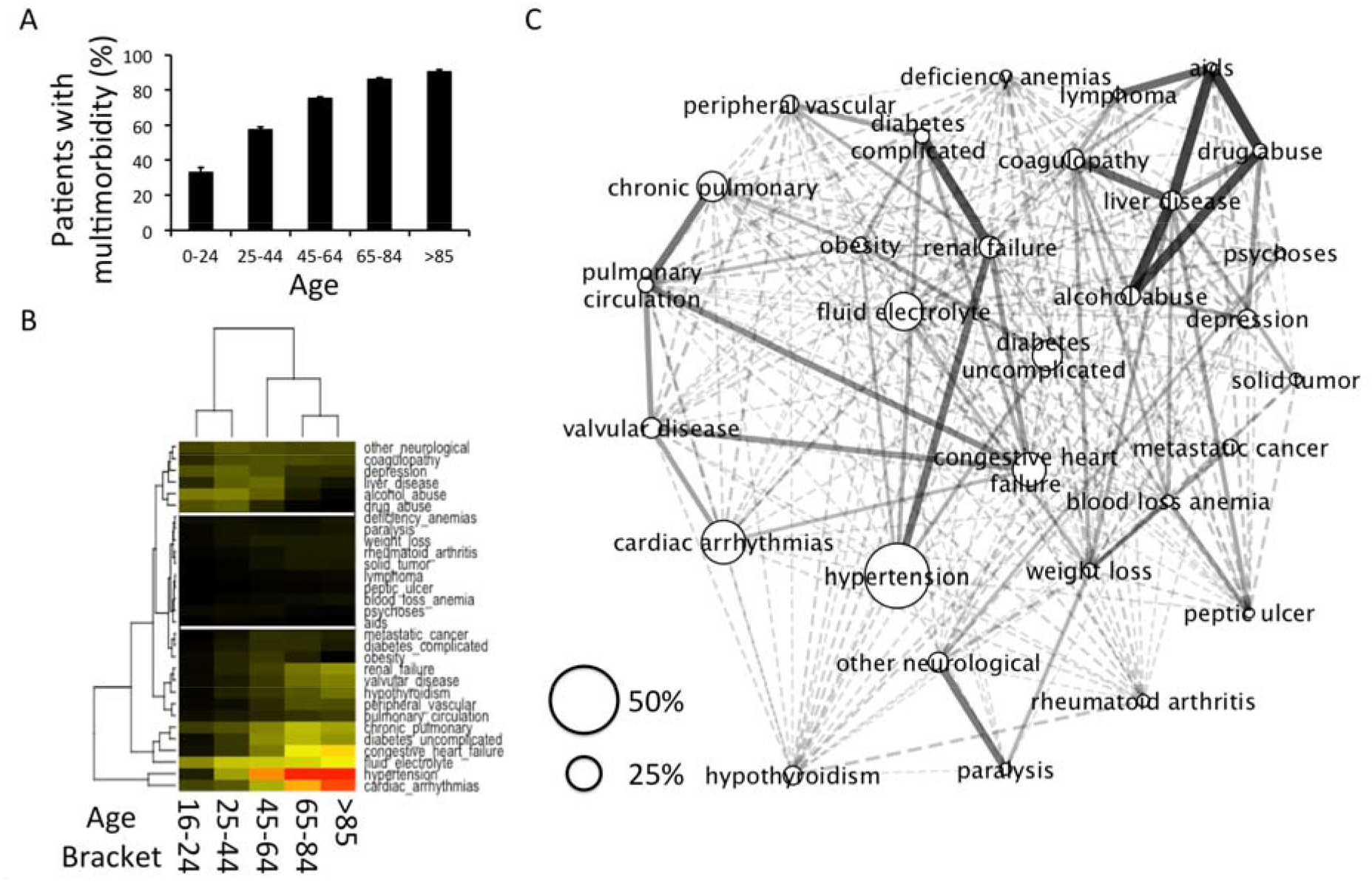
Heterogeneous morbidity profile in critical care patients. A: Percent of patients suffering from multimorbidity progressively increases with age. B: Disease composition is non-homogenous over each age bracket but organizes into the following patterns: (1) diseases that are more prevalent increase progressively with age, (2) constant with age, (3) have an earlier prevalence peak and become less common in older population. C: Network discovery of disease co-occurrence shows broad range of disease associations in the critical care study cohort. Note intuitive examples such as associated cardiopulmonary conditions, diabetic nephropathy, co-occurrence of alcohol abuse, liver disease and coagulopathy. Node size represents disease prevalence; edge width expresses relative risk for each disease pair.

### Identifying distinct multimorbidity endophenotypes

We hypothesized that with analysis of similarities at the individual patient level, we can identify subgroups of patients with distinct demographics and disease compositions. We also hypothesized that patients belonging to some of these subgroups will share vulnerability to sepsis and associated death. Using latent class analysis we identified six classes of patients, referred to as Class 1-6 from here onwards. The number of classes were defined by considering metrics of model fit, class size and interpretability. Classes were verified using descriptive statistics as well as simulations (see Supplementary material and Methods). Disease compositions paralleled some of the pattern suggested by the previous network discovery of the entire study cohort (Fig1C). Endophenotype class characteristics are summarized in Fig 2, Fig 3 and Supplementary Table S2.

**Figure 2:**
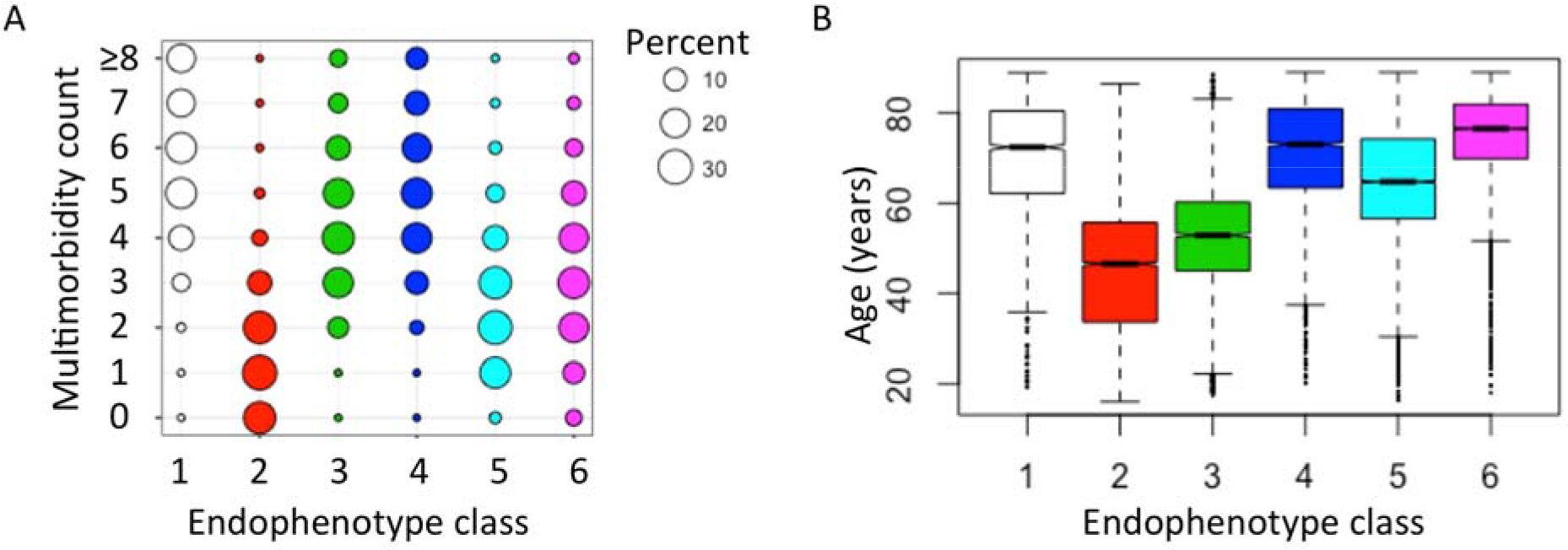
Class characteristics of the six endophenotypes identified in critical care cohort. A: Baloonplot summary of morbidity count for each endophenotype class. Highest morbidity burdens are class 1, 3, 4 and 6. B: Boxplot of age distribution in endophenotype classes.

**Figure 3:**
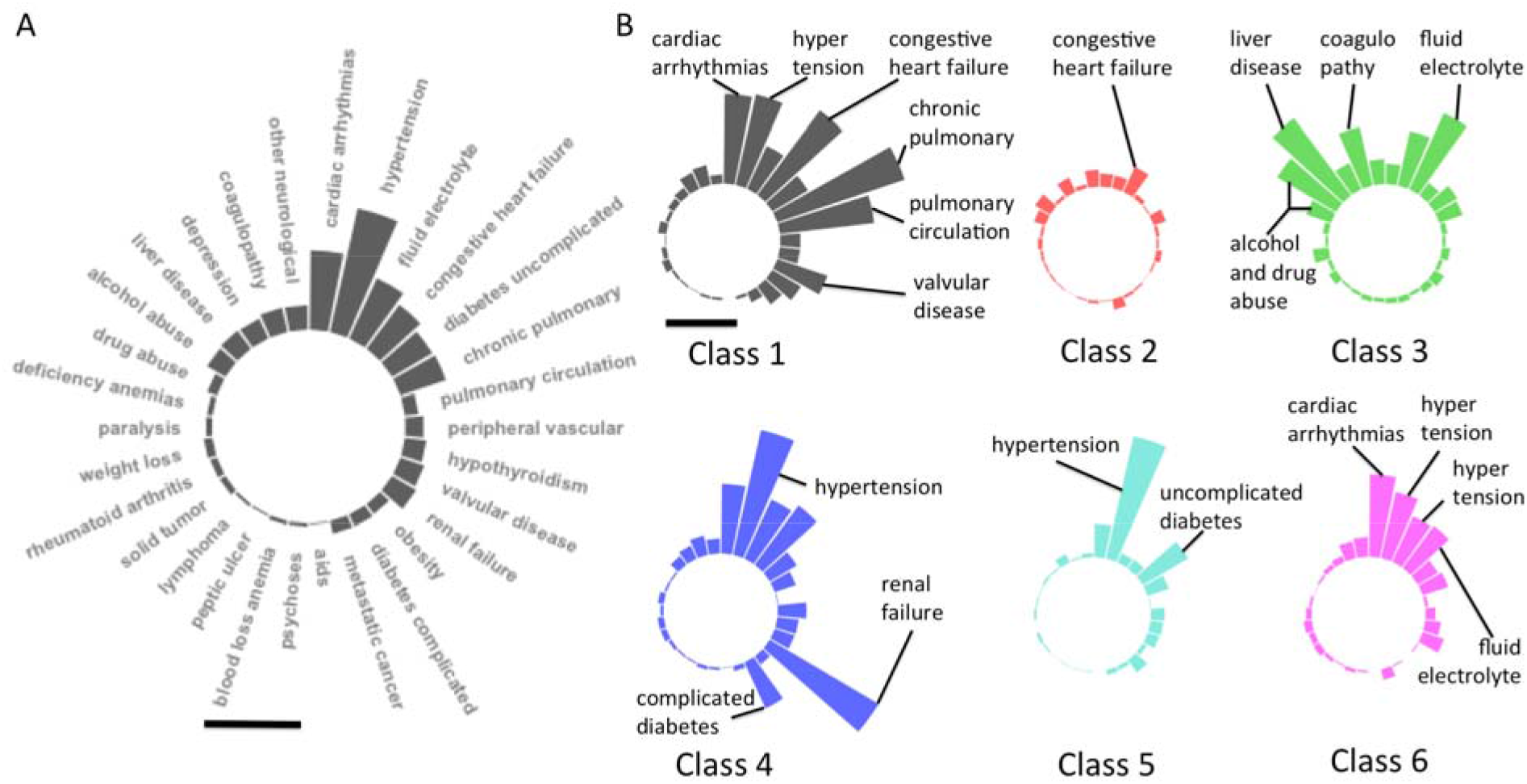
Distinct disease compositions in critical care endophenotypes. A: Prevalence of Elixhauser’s morbidity categories for the entire cohort of 36960 patients. Bar=50%. B: Circular barplot summary of disease composition of morbidity endophenotypes in critical care. Plot configuration is identical to “A”. Note the distinct patterns such as the complex cardiopulmonary profile in class 1, health consequences of addiction in class 3 and diabetic nephropathy with hypertension in class 4. Bar=50%.

In Class 1 we found high prevalence of cardiopulmonary conditions in older patients (mean age 72.3 ± 0.27) with the highest prevalence of chronic pulmonary diseases (93.86%). Class 2 had the youngest patients with the lowest point prevalence of any morbidity compared to the rest of the classes. Class 3 consisted of middle-aged patients (mean age of 52.25, 95% CI 51.85-52.65) with the high rates of depression (20.1%), alcohol abuse (47.75%), drug abuse (18.2%), liver failure (67%), and coagulopathy (41.81%). This class captured 70% of patients with liver disease from the entire critical care cohort of 36,960 patients. Class 4 and Class 5 both had high prevalence of hypertension, and a large proportion of patients suffered from renal failure (88,3%) and complicated diabetes (35,4%) in class 5. Class 6 was comprised of the oldest patients with high prevalence of cardiopulmonary problems similar to Class 1. Statistical testing with one-way ANOVA showed significant differences between all disease prevalence between classes (p<0.001) supporting the distinct disease composition of each endophenotype, the basis for identification of these subgroups.

### Multimorbidity endophenotypes vulnerable to sepsis and associated mortality

We next tested if multimorbidity endophenotypes shared vulnerability to sepsis and death. We first established the patterns of organ dysfunctions using Sequential Organ Failure Assessment (SOFA) score which entails a system-wise observations recorded during the first 24 hours of ICU stay (3). The system level assessment using the SOFA subscores for respiratory, cardiovascular, renal, hepatic, coagulation and central nervous systems paralleled the morbidity profile of each endophenotype class (Supplementary Fig S1). The highest SOFA scores were detected in Class 3 with closely followed by Class 4 risk scores (Fig 4A). OASIS risk assessment score for inpatient mortality showed higher values for classes 1, 3, 4 and 6 (Fig 4B). The actual rate of organ failure and sepsis were highest in Class 3 followed by Class 4 (Fig 4C). High mortality classes were Class 1, 3, 4 and 6. Mortality related to sepsis and organ dysfunction was highest in Class 3, almost fourfold the mortality rates in Classes 2 and 5 (Fig 4D).

**Figure 4:**
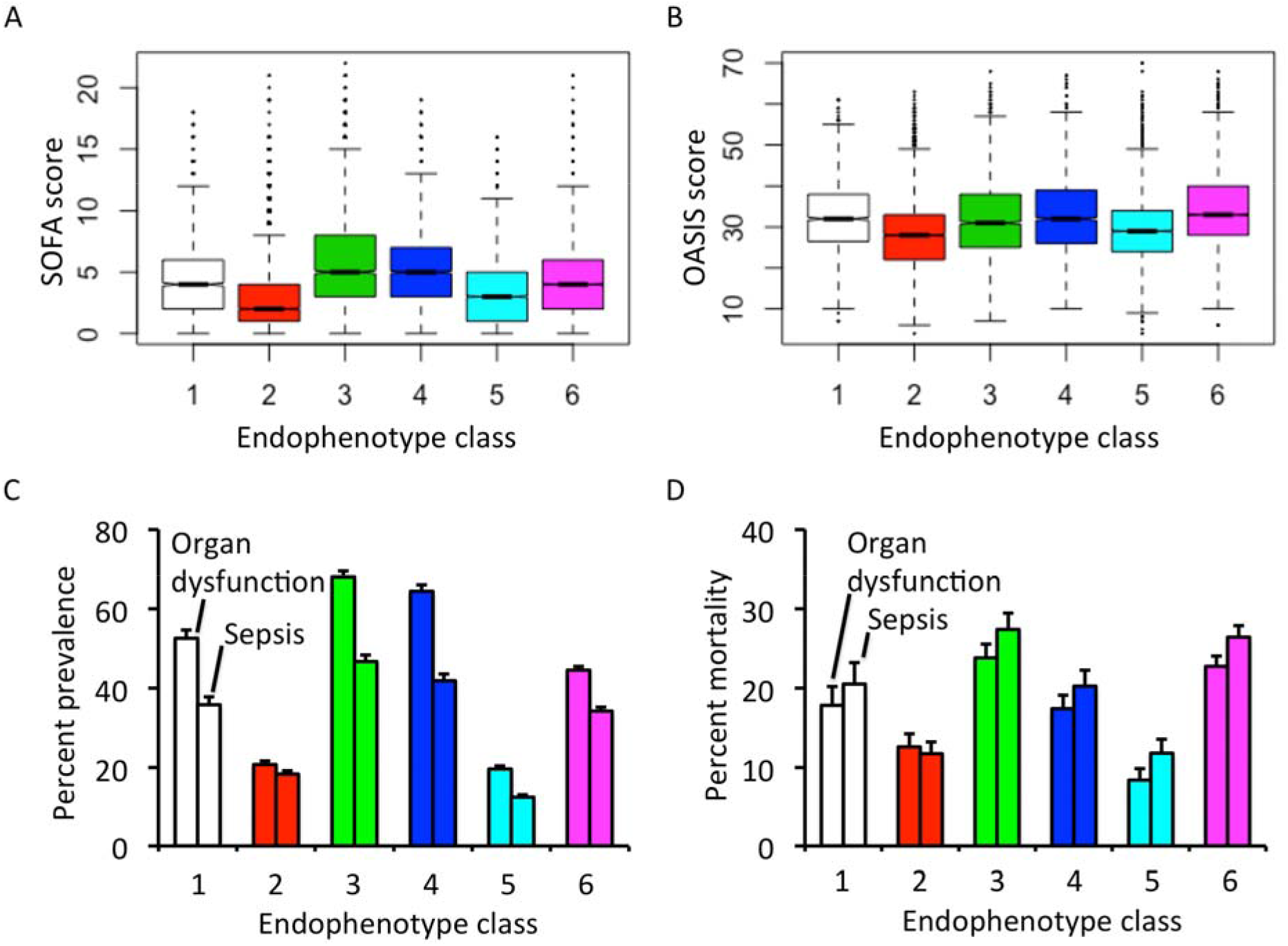
Morbidity endophenotypes vulnerable to organ dysfunction, sepsis and associated mortality. A: SOFA score shows most severe organ failure in class 3 closely followed by class 4. B: OASIS score shows similarly high mortality risk in classes 1, 3, 4 and 6. C and D: Prevalence of organ dysfunction, sepsis and associated mortality are highest in endophenotype 3. (all differences are significant with multiple testing, p<0.001)

### Disease co-occurrence in vulnerable multimorbidity endophenotypes

We next explored the co-occurrence of conditions for endophenotype classes 1, 3, 4 and 6 with high risk of adverse health outcomes (sepsis, organ dysfunction and death) by visualizing prevalence and pairwise associations in a disease network (Fig 5).

**Figure 5:**
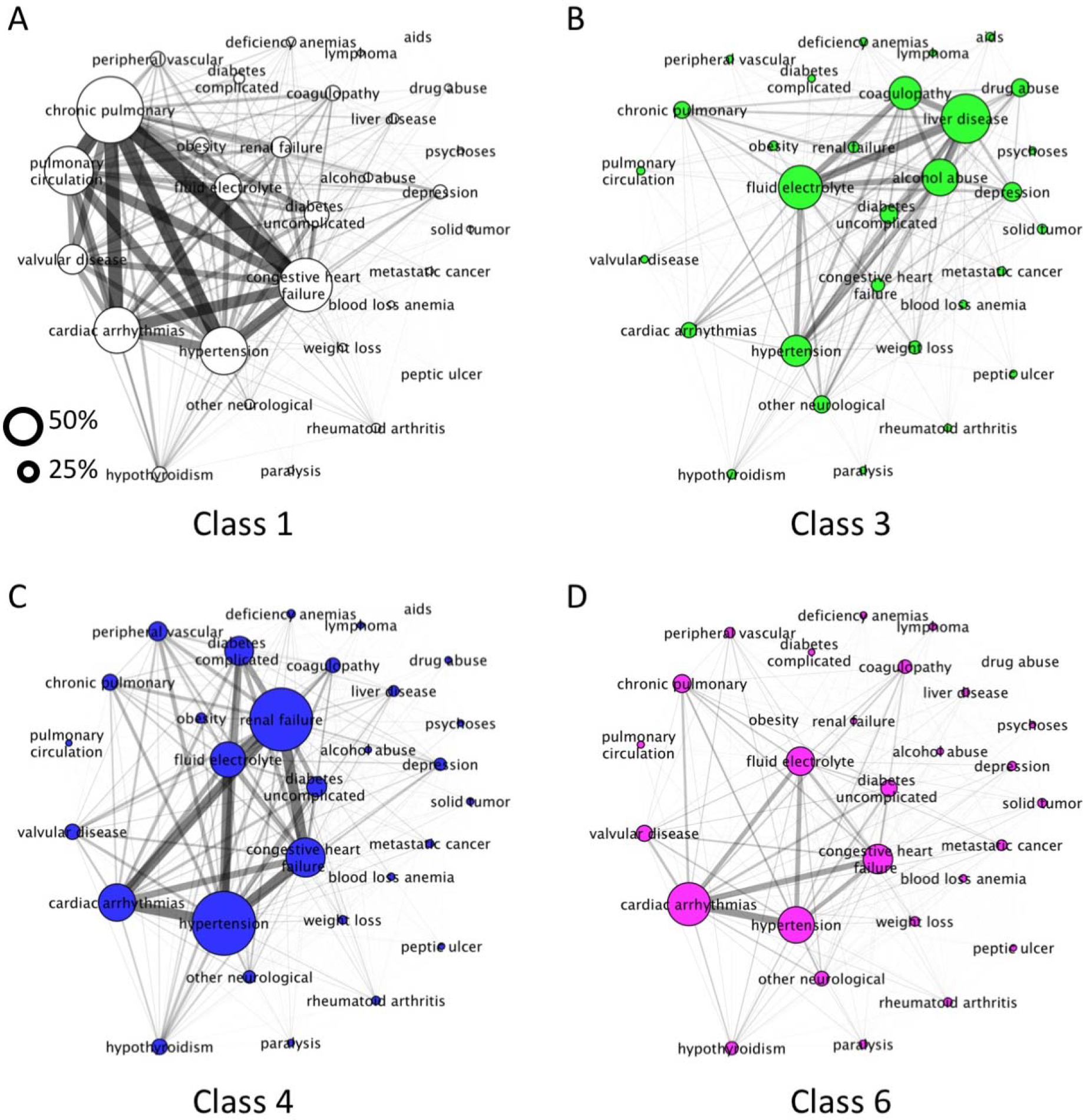
Network discovery of multimorbidity endophenotype classes with higher rates of adverse health outcomes (organ failure sepsis and death). A: Patients in Class 1 suffer from cardiopulmonary diseases. B: Class 3 has a high prevalence of health consequences of addiction. C: For patients in Class 4 the dominant morbidity profile is diabetic nephropathy and hypertension. D: Class 6 have suffer from cardiopulmonary comorbidities. Nodes represent disease prevalence, edges express the number of patients with the disease pair normalized to the entire class population.

Nodes represented the Elixhauser disease categories with size defined by disease prevalence within the subgroup and edge width was the number of patients with the disease pair normalized to the total number of patients in the endophenotype class (Fig 3). Relative risk and/or Pearson’s correlation were not used in this portion of the analysis as these techniques tend to render inaccurate results for extremes of prevalence potentially resulting in under/over-estimated associations (25). Network structure confirmed disease patterns suggested by our analysis of disease prevalence in each of the endophenotypes. The high mortality group consisting of younger patients and suggested a pattern of addiction-associated conditions (Class 3) given the pairwise association between liver disease, coagulation disorders, alcohol excess, drug abuse and depression. In further Classes the networks analysis showed associations between cardiovascular-respiratory conditions (Class 1), hypertensive-renal-diabetic with end organ complications (Class 4), and cardiopulmonary problems (Class 6).

Groups with lower mortality (5.31-6.02%) were either younger patients (99% below 65 years, Class 2) with very low disease burden or elderly with significantly higher proportion of non-emergency admissions (35.7%, Class 5) than the rest of the cohort. This Class suffered from combination hypertensive-diabetes without end organ complications (Supplementary Fig 2).

## Discussion

Our study identifies multiple clinical subgroups in a critical care cohort with high rates of organ dysfunction, sepsis and associated mortality. In addition to the expected phenotypes of multimorbid elderly with high mortality we also find a group of younger patients suffering from health consequences of addiction and the highest rates of sepsis and organ dysfunction. Although the association of liver failure with poor outcomes in sepsis is well established, the multimorbidity configuration it occurs in is less appreciated. These findings support the shift away from the single disease model in healthcare to a more holistic construct, putting emphasis on the consideration of multimorbidity composition in clinical decision making.

As expected, we found increasing prevalence of multimorbidity with age in line with previous population studies (9). However several of our identified endophenotypes demonstrated distinct morbidity composition, high rates of sepsis and associated mortality. The disease combinations within these subgroups (Class 1, 3, 4 and 6) were in line with previous observations and can also be interpreted along the lines of shared risk factors and patho-mechanisms. For example, chronic pulmonary disease and congestive heart failure co-occur with the highest prevalence in Class 1, this may be explained by the concept of the “cardio-pulmonary continuum” (26). Class 4 corresponds to a patient group suffering from diabetic nephropathy and hypertension while Class 6 would represents elderly with cardiovascular diseases and higher rate of neurological conditions.

The most vulnerable group to adverse health outcomes consisted of younger/middle-aged population who suffered from high prevalence of alcohol and/or drug addiction, associated with liver disease and coagulopathy. While liver disease is well recognized as a risk of sepsis as well as poor outcomes, to the best of our knowledge our results are the first to 1) identify complex multimorbidity phenotypes incorporating chronic liver disease 2) compare it with other disease configurations in terms of sepsis prevalence and mortality. The demographics and disease composition of this high-risk group paralleled the data from US Center for Disease Control and Prevention that showed peak percentage of alcohol consumption in the age bracket of 25-44 and 77% of deaths from alcoholic liver disease under the age of 65 (27). Population studies covering 175 million hospital discharges demonstrated that the 4.5 million patients suffering from liver cirrhosis were twice as likely to die while in hospital, have sepsis as the reason for admission and die of sepsis (28, 29). Bacterial infections are present in up to 30% of admissions with liver failure (30). The liver has widespread functions in responding to sepsis such as synthesis of proteins for immune, coagulation and metabolic functions as well as scavenging of endotoxins and bacteria (31–33). Defects in the sub-mechanisms of the immune system such as macrophage Fcγ receptor mediated clearance, complement system, monocyte HLA-DR expression, neutrophil phagocytosis and natural killing. There is also element of hyperactivity in cirrhosis consisting of abnormally upregulated TNF-α and IL6 in response to infection. This effect was studied in animal models of ischemia reperfusion injury to the liver and experimental bacteremia, where greater interleukin and TNF-α production and consequent lung inflammation was detected compared to sham animals (38). This paralells the higher rate of organ failure traceable in our own findings with the highest SOFA score in class 3 of our endophenotypes (Fig 4A, Supplementary Fig S1).

Conventionally the recruitment into clinical trials in sepsis have been based on abnormal physiological parameters implying infection as cause for critical illness. Such recruitment strategies inevitably capture a heterogenous patient group where the morbidity profile would associate with alterations in the biological response to sepsis. Interventions often studied by clinical trials included modifiers of inflammatory response in sepsis such as anti-TNF, anti-IL1-Ra, anti-LPS, corticosteroids, IV immunoglobulins and activated protein C (34). On the other hand conditions such as coagulopathy, chronic liver disease (35), diabetes (36) with end organ complications are characterized by alterations in these target mechanisms. As described above, excessive levels of inflammatory mediators produced by the impaired/cirrhotic liver may underlie the failure of trial treatments in subgroup (28, 37). In diabetes multiple components of sepsis immune response are effected, for example by neutrophil dysfunction related to hyperglycemia, adaptive immunity and IL6 signaling (36). Consequently, the proposed multimorbidity endophenotypes in our study, in particular groups with liver disease or complicated diabetes may represent higher rates of non-responders in clinical trials.

It is well recognized that managing sepsis patients with liver failure is challenging and requires an individualized form of goal-directed therapy (39). Early diagnosis and treatment are essential. The early identification of latent classes may therefore help decision-making regarding prophylactic antibiotics, consideration of early interventions, and lower threshold for goal directed therapy to improve outcomes in sepsis.

One potential limitation of this study is that it’s based on data from a single center, therefore composition of catchment population, departmental protocols, resources and staffing characteristics are potential limits to the generalizability of our results (40). Further, it is difficult to find external validation to our results because MIMIC3 is a unique dataset in the high-level levels of clinical detail it provides as well as patient volume. Another limitation is that, we analyze a cross section of the population and therefore cannot examine causality links between endophenotypes and vulnerability to sepsis and organ dysfunction. For the same reason we have no information on time of diagnosis to allow us to distinguish between what is pre-existing medical condition versus new diagnosis. To address this issue drug prescription records were incorporated to filter out diseases that are the reason for the hospital stay versus those that are not. Finally, our study is limited to patients that survive long enough or are deemed appropriate for intensive care. Therefore, mortality numbers very likely underestimate true figures. Nevertheless our study uses real world hospital data and provides a description of the population in the first 24 hours in critical care, a very relevant time frame of management. This provides critical information on risk of sepsis and associate mortality.

## Conclusions

The increasing prevalence of multimorbidity creates complexity in medical diagnostics and treatment decisions. Our study examines the relevance of multimorbidity in sepsis, one of the highly relevant conditions in critical care. We identify several patient groups susceptible to adverse health outcomes. The patient group most vulnerable to sepsis, organ failure and mortality was a less anticipated class of patients suffering from health consequences of addiction. While this work is hypothesis generating, the findings support the shift away from current single-disease model and promote an endophenotype specific approach in medical training, treatment and trial design.

## Supporting information

## Author contributions

ZZ and NG conceived the study, ZZ: carried our data analysis, ZZ, AL, MC, NG: wrote the paper

## Acknowledgements

The authors would like to thank Ling Chen for her help with parts of the analysis, Erik Drysdale and Lauren Erdman for their helpful discussions on the manuscript.

## Data Availability

MIMIC 3 database is available at: https://mimic.physionet.org/about/mimic/ Codes used to generate data tables are accessible at: https://github.com/MIT-LCP/mimic-code

