## Supplementary material for "“Multimorbidity states with high sepsis-related deaths: a data-driven analysis in critical care”"

**Supplementary methods**

*Database*

MIMIC 3 is a publicly accessible prospectively gathered database over a period of 7 years. It consists of 58,976 unique hospital admissions and 61,051 unique critical care admissions for a total of 46,428 patients with unique subject identifiers. Data were merged through unique identifiers using *sql*; yielding a total of 38,557 adult patients who were admitted for the first time to critical care. Following a complete data analysis 2,167 (5.6%) patients were excluded for missing data. This process is summarized in supplementary Fig S1. The population characteristics of the remaining 36,390 adults are summarized in Supplementary Table 1.

*Latent class analysis (LCA)*

LCA is a well-established statistical method used extensively in defining subtypes of asthma (1), acute respiratory distress syndrome (2) or Alzheimer’s disease (3). Conventional regression techniques set out to find associations between known variables and an outcome. On the other hand LCA defines subgroups as a combination of measured variables and relates them to a set of unobserved variables using a subset of structural equation modeling. The resulting classes are mutually exclusive and patients within each class share clinical similarities. We favoured this approach due to its capability to incorporate unobserved variables in contrast to other clustering techniques, which are often based on only observed variables. We applied LCA to establish endophenotypes using the input variables available to physicians at the time of admission to critical care; specifically, age, sex, type of admission, and morbidities. We used the Bayesian Information Criterion (BIC) and Akaike Information Criterion (AIC) to objectively assess model fit. This technique balances model complexity against predictive accuracy in order to determine the optimal number of latent classes. Our choice of class number was guided by the combination of the following: 1) lowest possible AIC and/or BIC value, 2) smallest class size at or above 5% of the core dataset (36390 patients) with interpretable class as an additional consideration (3, 4). We verified the preferred homogeneity within classes and heterogeneity between classes by Chi square and t-test for categorical and continuous variables respectively. We also assessed the ability for input variables to predict class membership and carried out a regression analysis with 10-fold cross validation as described previously (5, 6). We created dichotomized labels for 1) subclass of interest 2) all other subclasses and trained a logistic regression model with the same input variables used in identifying the latent classes using the LCA. Model performance was assessed using area under the receiver operating curve, which plots model performance (sensitivity versus specificity) across a range of thresholds differentiating two groups (Supplementary figure 3, Supplementary table 3).

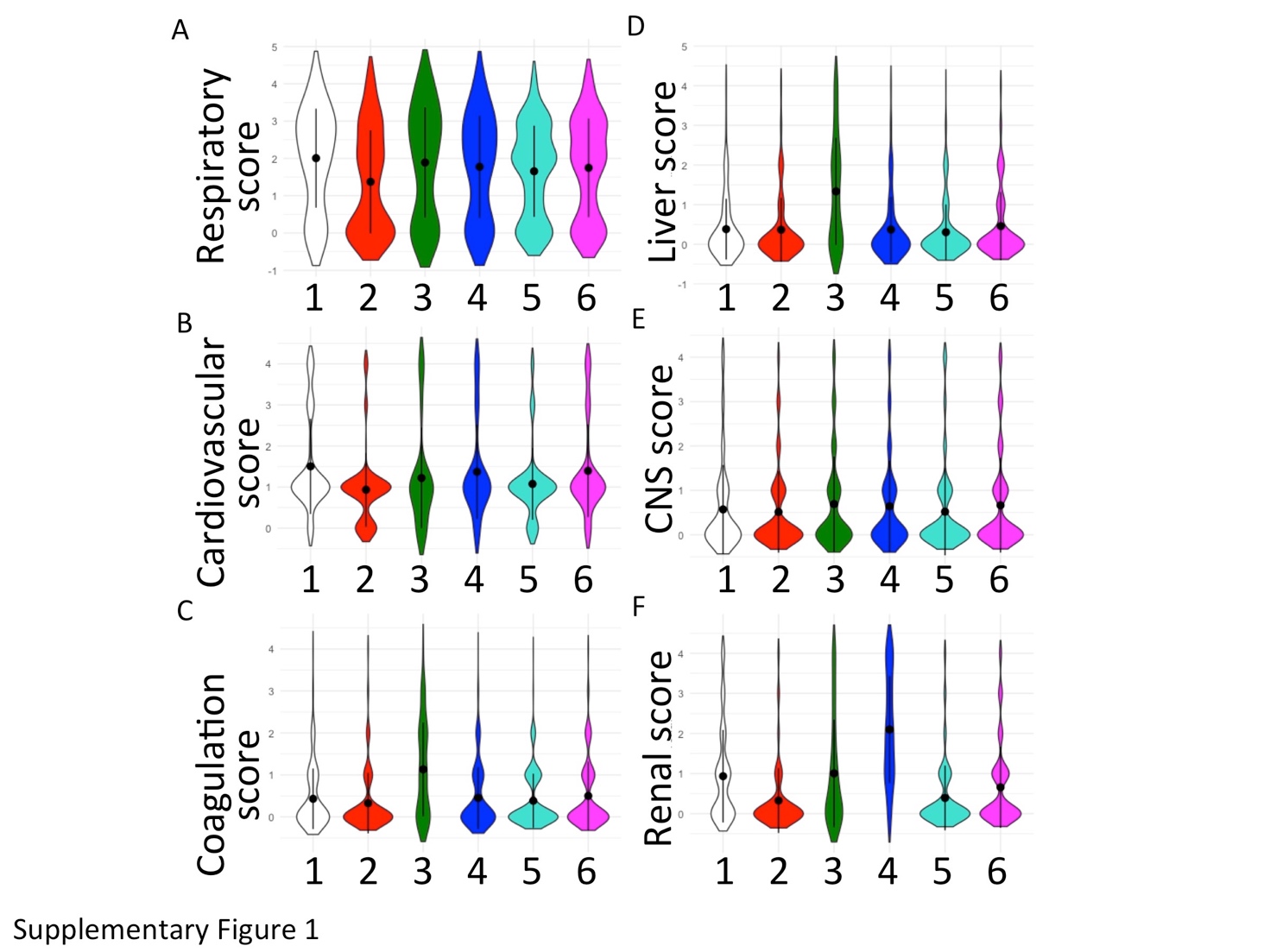

**Supplementary Figure S1:** Violin plot summary of organ systems impairment in the multimorbidity endophenotype classes.

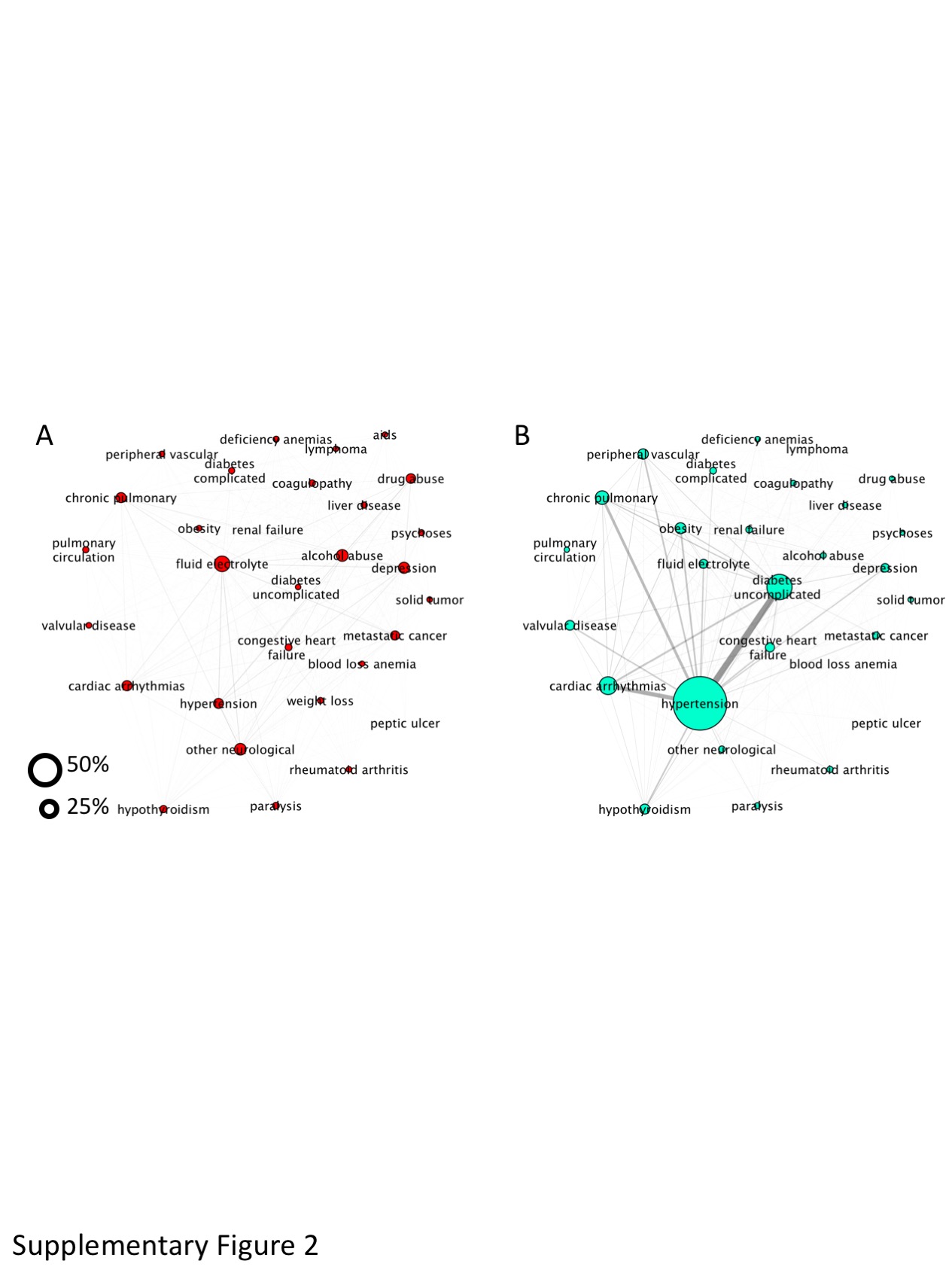

**Supplementary Figure S2:** Network summary of multimorbidity endophenotype classes with lower rates of sepsis and death.

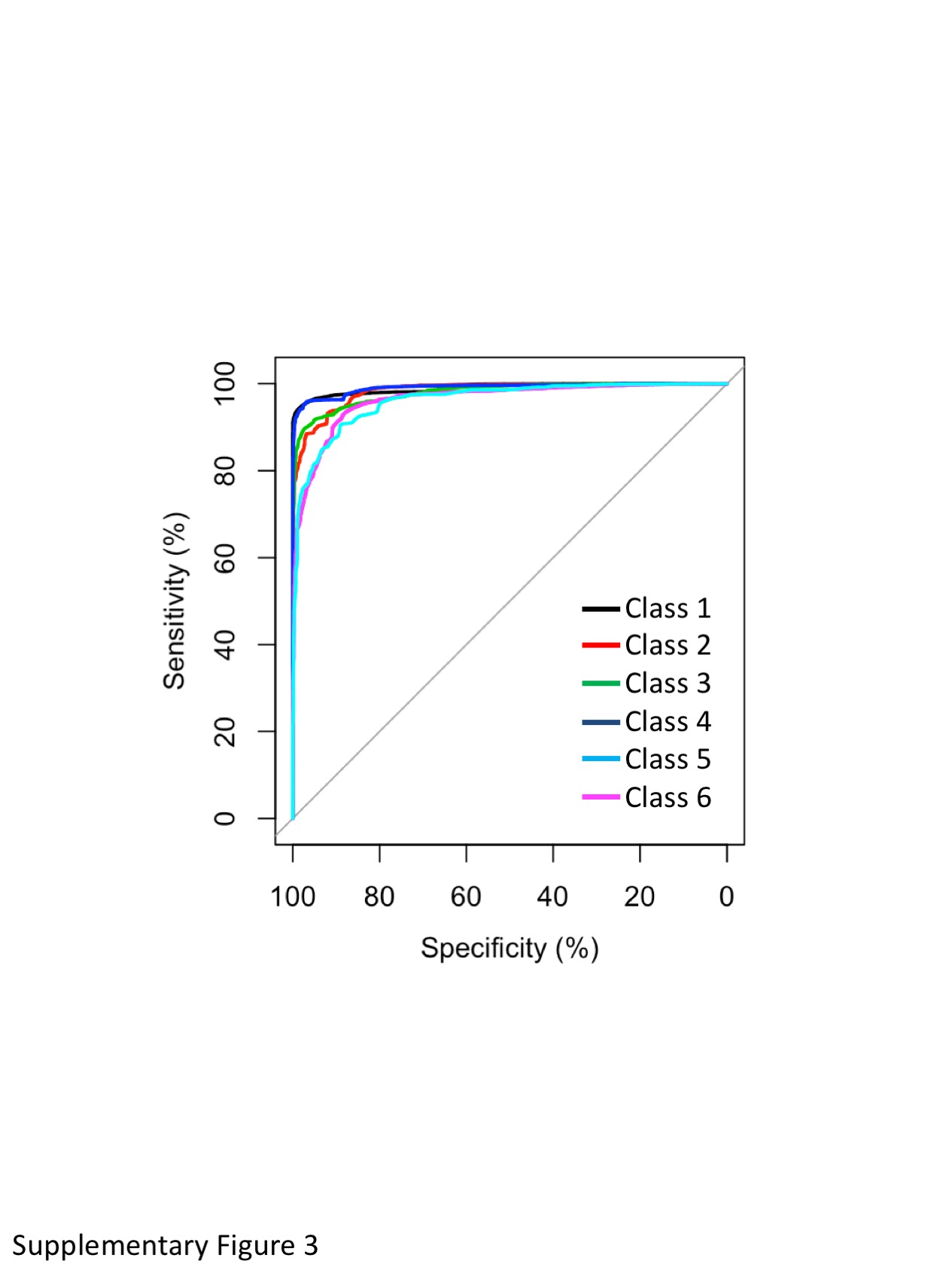

**Supplementary Figure S3:** Graph summary of ROC curves depicting predictive performance of LCA input variables at differentiating a given class from the remaining classes.
