## Supplementary material for "“Multimorbidity states with high sepsis-related deaths: a data-driven analysis in critical care”"

**Supplementary Table S1: Demographics and outcomes summary of study cohort**

|  | **Number of  patients (%)** | **Median morbidity count (IQR)** | **Percent (95% CI)  with multimorbidity** | **SOFA score**  **(95% CI)** | **LOS ICU**  **(95% CI)** | **LOS Hospital**  **(95% CI)** | **Percent Mortality**  **(95% CI)** |
| --- | --- | --- | --- | --- | --- | --- | --- |
| **All patients** | 36390 (100%) | 3（2-4） | 77.27%  （76.84%, 77.70%) | 3.99  (3.96, 4.02) | 4.12  (4.06, 4.18) | 10.06  (9.95, 10.17) | 10.9  (10.6, 11.3) |
| **Sex** |  |  |  |  |  |  |  |
| male | 21052 (57.85%) | 3 (2-4) | 76%  (75.42%, 76.57%) | 4.13  (4.09, 4.17) | 4.07  (4, 4.17) | 10.06  (9.91, 10.21) | 10.3  (9.95, 10.8) |
| female | 15338 (42.15%) | 3 (2-4) | 79.02%  (78.37%, 79.66%) | 3.79  (3.74, 3.84) | 4.17  (4.07, 4.27) | 10.06  (9.89, 10.22) | 11.7  (11.2, 12.2) |
|  | **Age** |  |  |  |  |  |  |
| 0-24 | 1220 (3.35%) | 1 (0-2) | 33.11%  （20.53%, 35.80%） | 2.63  (2.48, 2.77) | 3.4  (3.12, 3.69) | 8.12  (7.56, 8.67) | 4.4  (3.3, 5.8) |
| 25-44 | 4563 (12.54%) | 2 (1-3) | 57.75%  (56.31%, 59.17%) | 3.29  (3.2, 3.38) | 4.12  (3.92, 4.32) | 10.1  (9.71, 10.47) | 6.02  (5.4, 6.8) |
| 45-64 | 12831 (35.26%) | 3 (2-4) | 75.22%  (74.46%, 75.96%) | 3.92  (3.87, 3.98) | 4.07  (3.97, 4.18) | 10.4  (10.2, 10.6) | 8.5  (8.1, 9) |
| 65-84 | 15691 (43.12%) | 3 (2-5) | 86.31%  (85.76%, 86.84%) | 4.3  (4.26, 4.35) | 4.25  (4.16, 4.35) | 10.1  (9.9, 10.2) | 13.5  (13, 14) |
| 85-95 | 2085 (5.73%) | 4 (2-5) | 90.46%  (89.12%, 91.64%) | 4.3  (4.21, 4.46) | 3.87  (3.64, 4.1) | 9.12  (8.79, 9.45) | 20.8  (19.1,22.6) |
| **No. of disorders** |  |  |  |  |  |  |  |
| 0 | 2669 (7.33%) | **-** | **-** | 2.53  (2.44, 2.62) | 3.1  (2.89, 3.29) | 7.56  (7.21, 7.92) | 5.8  (4.99, 6.81) |
| 1 | 5602 (15.39%) | **-** | **-** | 2.95  (2.88, 3.02) | 3.3  (3.16, 3.45) | 8.18  (7.92, 8.45) | 7.27  (6.6, 7.99) |
| 2 | 7410 (20.36%) | **-** | **-** | 3.54  (3.48. 3.6) | 3.76  (3.62, 3.89) | 9.2  (8.95, 9.44) | 8.99  (8.35, 9.67) |
| 3 | 7088 (19.84%) | **-** | **-** | 4  (3.93, 4.07) | 4  (3.88, 4.14) | 9.86  (9.63, 10.1) | 11.1  (10.3, 11.8) |
| 4 | 5477 (15.05%) | **-** | **-** | 4.5  (4.42, 4.59) | 4.32  (4.16, 4.48) | 10.57  (10.3, 10.9) | 12.2  (11.4, 13.2) |
| 5 | 3772 (10.37%) | **-** | **-** | 4.86  (4.76, 4.97) | 4.86  (4.64, 5.08) | 11.71  (11.35, 12.1) | 13.7  (12.6, 14.8) |
| 6 | 2218 (6.1%) | **-** | **-** | 5.23  (5.09, 5.37) | 5.22  (4.93, 5.52) | 12.54  (12, 13.1) | 16.2  (14.7, 17.8) |
| 7 | 1213 (3.33%) | **-** | **-** | 5.68  (5.49, 5.87) | 5.82  (5.41, 6.24) | 14.16  (13.4, 14.92) | 17.6  (15.5, 19.8) |
| >8 | 941 (2.59%) | **-** | **-** | 6.11  (5.49, 5.87) | 6.74  (6.22, 7.26) | 15.75  (14.9, 16.6) | 21.8  (19.2, 24.6) |
| **Admission type** |  |  |  |  |  |  |  |
| elective | 5939 (16.32%) | 3 (1-4) | 74.61%  (73.49%, 75.70%) | 4.04  (3.97, 4.11) | 3.3  (3.17, 3.43) | 9.22  (8.96, 9.49) | 2.63  (2.24, 3.07) |
| non-elective | 30451 (83.68%) | 3 (2-4) | 77.79%  (77.32%, 78.25%) | 3.98  (3.94, 4.01) | 4.28  (4.21 4.35) | 10.22  (10.1, 10.34) | 12.6  (12.2, 12.9) |
