## Supplementary material for "“Multimorbidity states with high sepsis-related deaths: a data-driven analysis in critical care”"

**Supplementary Table S2:** summary of morbidity composition of endophenotypes.

| **Condition** | **Class 1** | **Class 2** | **Class 3** | **Class 4** | **Class 5** | **Class 6** |
| --- | --- | --- | --- | --- | --- | --- |
| Number of patients | 2216 | 8540 | 3583 | 3428 | 9019 | 9604 |
| congestive heart failure | 73.42 | 3.43 | 11.14 | 51.46 | 7.81 | 36.53 |
| cardiac arrhythmias | 62.41 | 9.34 | 14.37 | 48.57 | 23.12 | 57.40 |
| valvular disease | 36.91 | 1.35 | 2.71 | 14.67 | 8.56 | 15.75 |
| pulmonary circulation | 66.29 | 2.35 | 3.85 | 0.70 | 0.62 | 2.11 |
| peripheral vascular | 14.17 | 1.53 | 2.40 | 20.13 | 9.87 | 7.00 |
| hypertension | 64.67 | 9.16 | 38.93 | 89.47 | 87.70 | 47.14 |
| paralysis | 0.68 | 3.20 | 1.95 | 2.16 | 1.76 | 3.86 |
| other neurological | 6.41 | 12.06 | 17.44 | 10.47 | 2.96 | 13.61 |
| chronic pulmonary | 93.86 | 9.18 | 17.22 | 15.61 | 15.67 | 18.78 |
| diabetes uncomplicated | 28.88 | 1.02 | 18.64 | 21.94 | 36.82 | 16.51 |
| diabetes complicated | 7.67 | 2.14 | 2.46 | 35.41 | 3.03 | 1.02 |
| hypothyroidism | 14.21 | 4.16 | 5.78 | 15.11 | 9.17 | 11.06 |
| renal failure | 23.06 | 0.29 | 7.98 | 88.30 | 3.28 | 0.99 |
| liver disease | 6.45 | 2.39 | 67.01 | 7.56 | 1.19 | 3.29 |
| peptic ulcer | 0.54 | 0.27 | 2.29 | 1.11 | 0.58 | 0.93 |
| aids | 0.09 | 0.71 | 3.80 | 0.47 | 0.00 | 0.00 |
| lymphoma | 1.26 | 0.88 | 1.67 | 1.28 | 0.29 | 2.50 |
| metastatic cancer | 2.21 | 7.45 | 4.47 | 3.56 | 3.66 | 7.47 |
| solid tumor | 1.90 | 1.30 | 5.89 | 2.36 | 1.20 | 4.83 |
| rheumatoid arthritis | 5.32 | 1.41 | 2.65 | 4.29 | 2.34 | 4.27 |
| coagulopathy | 14.35 | 2.87 | 41.81 | 13.80 | 1.21 | 11.96 |
| obesity | 15.88 | 1.38 | 5.30 | 7.15 | 10.93 | 0.02 |
| weight loss | 3.47 | 1.99 | 11.16 | 4.38 | 0.00 | 5.76 |
| fluid electrolyte | 32.99 | 19.29 | 58.39 | 45.45 | 7.15 | 34.96 |
| blood loss anemia | 1.76 | 1.11 | 3.77 | 2.60 | 0.49 | 2.56 |
| deficiency anemias | 5.91 | 1.99 | 3.96 | 4.14 | 1.03 | 1.88 |
| alcohol abuse | 4.96 | 12.48 | 47.75 | 1.28 | 1.67 | 1.51 |
| drug abuse | 2.62 | 8.43 | 18.22 | 1.55 | 0.62 | 0.36 |
| psychoses | 1.85 | 2.11 | 2.85 | 1.78 | 0.63 | 1.31 |
| depression | 12.50 | 11.17 | 20.12 | 10.04 | 6.43 | 4.65 |
