## Supplementary material for "“Multimorbidity states with high sepsis-related deaths: a data-driven analysis in critical care”"

**Supplementary table S3.** Table summary of classifier performance (assessed using area under the receiver operating curve, AUC) at predicting endotype class based on the input variables used in the LCA (age, sex, type of admission, and morbidities).

| Class | AUC | 95% CI |
| --- | --- | --- |
| 1 | 0.9907 | 0.9899-0.9915 |
| 2 | 0.9833 | 0.9823-0.9843 |
| 3 | 0.9799 | 0.9785-0.9812 |
| 4 | 0.9909 | 0.9902-0.9917 |
| 5 | 0.9612 | 0.9592-0.9631 |
| 6 | 0.963 | 0.9611-0.9649 |
